# An untargeted metabolomic approach to identify antiviral defense mechanisms in memory leukocytes secreting *in vitro* IgG anti-SARS-Cov-2

**DOI:** 10.1101/2021.07.20.453042

**Authors:** Gevi Federica, Fanelli Giuseppina, Lelli Veronica, Zarletti Gianpaolo, Tiberi Massimo, De Molfetta Veronica, Scapigliati Giuseppe, Timperio Anna Maria

## Abstract

Available knowledge shows that individuals infected by SARS-CoV-2 undergo an altered metabolic state in multiple organs. Metabolic activities are directly involved in modulating the immune responses against infectious diseases, yet our understanding remains limited on how host metabolism relates with inflammatory responses. To better elucidate the underlying biochemistry of leukocytes response, we focused our analysis on the possible relationships between SARS-CoV-2 post-infection stages and distinct metabolic pathways. Indeed, in cultures of peripheral blood mononuclear cells (PBMC, n=48) obtained 60-90 days after infection and showing in vitro IgG antibody memory for spike-S1 antigen (n=19), we observed a significant altered metabolism of tryptophan and urea cycle pathways. This work for the first time identifies metabolic routes in cell metabolism possibly related to later stages of immune defense against SARS-Cov-2 infection, namely when circulating antibodies may be absent, but an antibody memory is present. The results suggest a reprogramming of leukocyte metabolism after viral pathogenesis through activation of specific amino acid pathways possibly related to protective immunity against SARS-CoV-2.

## Introduction

Covid-19 is the acute illness caused by SARS-CoV-2 with initial clinical symptoms such as cough, fever, malaise, headache, and anosmia^1^. Being a new form of virus, pathogenesis of the disease is not fully understood, contrasted by vaccination policies and potential drugs developed^2^. The viral infection stimulates immune responses, also directed against the spike protein (S1 protein) present on the surface of SARS-Cov-2, being a ligand that binds to ACE2 receptor in host cells. After entry into cells the corona viruses (CoV) activate aryl hydrocarbon receptors (AhRs) through an indoleamine 2,3-dioxygenase (IDO1)-independent mechanism. This leads to an unregulated up-stimulation of downstream effectors among which the modulation of proinflammatory cytokine gene expression and in particular of interleukin 1*β* (IL-1*β*), IL-6, and TNF-*α*, consistent with the role for AhR activation following viral infection^3^. When SARS-Cov-2 infection persists, inflammation activates IDO1 by massively overexpressing proinflammatory cytokines, a deleterious process known as cytokine storm. The resulting cytokine storm diffuses to other organs, leading to multiorgan damage. Secretion of such cytokines and chemokines attracts immune cells, notably monocytes and T lymphocytes, from the blood into the infected site^4^. In the respiratory tract, a recruitment of immune cells and lymphocytes from the blood might explain the lymphopenia and increased neutrophil–lymphocyte ratio seen in around 80% of patients with SARS-CoV-2 infection, and after contact with viral antigens most of the effector T cells undergo apoptosis. Next, a pool of memory T cells is formed to fight reinfection. In the case of a subsequent infection, CD4+ memory T cells become restimulated, activate B cells and other immune cells by producing cytokines, with the help of CD8+ memory T cells that kill virus-infected cells^5,6^. Among the host responses, changes in cell metabolic processes play a pivotal role, and multiple approaches are currently used to identify key pathways involved in SARS-CoV-2 infection. Indeed, during targeted and untargeted metabolomics analyses, altered tryptophan metabolism involved in the kynurenine regulating inflammatory and immunity pathways have been identified^7^. Accordingly, 77 metabolites including amino acids, lipids, polyamines, sugars, as well as their derivatives have been discovered to be altered in the plasma of Covid-19 patients with severe symptoms, with respect to those showing mild symptoms^8^. Moreover, also cytosine (reflecting viral load), kynurenine (reflecting host inflammatory response), nicotinuric acid, and multiple short chain acylcarnitines (energy metabolism) were altered, in agreement with a severe outcome of the pathology, as already reported^9^.

Despite an increasing knowledge on metabolic changes in Covid-19 patients, little attention has been paid to post-infection stages (>60 days) where immune memory becomes responsible to protect against possible SARS-CoV-2 reinfection^10^. Indeed, as for other viral pathologies such as severe acute respiratory syndrome (SARS) and Middle East respiratory syndrome (MERS), there is a concern about Covid-2 effects. These effects may have longterm consequences and, importantly, the criteria for “recovery” are still ill-defined^11^ and mainly focused on the reduction of respiratory symptoms. In this respect, a knowledge on possible metabolic pathways linked to leukocytes and related to later stages of antiviral defences may be of importance.

To this regard we performed metabolomic profiling coupled with multivariate statistics analysis obtained from 43 cell cultures of peripheral blood mononuclear cells (PBMC), 19 of which displayed an *in vitro* IgG memory for spike-S1 antigen 60-90 days after infection determined by a cell-ELISA assay^12^. The obtained results suggest involvement of amino acid pathways, and are discussed as a possible protective mechanism over time against SARS-Cov-2.

## Material and method

### Subjects Recruitment

43 subjects (24 males and 19 females,dataset) of different ages (ranging from 30 to 81 years) undergoing SARS-Cov-2 serological analysis (Centro Polispecialistico Giovanni Paolo I, Viterbo, I) were enrolled in this study from October 2020 to March 2021. All participants provided informed written consent to participate in the research project, and the study was approved by the Regional Ethical Board in Ospedale L. Spallanzani, Roma, (number 169, approval 22/07/2020) and, in accordance with the Helsinki Declaration, written informed consent was obtained from all subjects. Supplementary Table 1 shows demographic and clinical characteristics of the recruited patients in the whole sample and by age group.

### Cell-ELISA and PBMC cultures

Determination of in vitro IgG B cell memory for spike-S1 virus protein and of specific IgG were performed by Cell-ELISA in PBMC, from all donors as previously described without modifications^12^, employing spike-S1 coated wells from a commercial source, (details in datasets), experimental outputs were net absorbance values (A450 nm) with background subtracted. Parallel cultures of PBMC employed in Cell-ELISA were incubated in microplates (Costar, USA) without stimulants at 106/ml at 37°C for 48 h in 100 ul/well of RPMI medium containing 10% FCS and antibiotics, then centrifuged at 500xg, and pelleted cells immediately employed for metabolomic analysis.

### Metabolomic analysis and data processing

Each sample was added to 1000 μl of a chloroform/methanol/water (1:3:1 ratio) solvent mixture stored at −20 °C. The tubes were mixed for 30 min and subsequently centrifuged at 1000 × g for 1 min at 4 °C, before being transferred to −20 °C for 2–8 h. The solutions were then centrifuged for 15 min at 15,000×g and were dried to obtain visible pellets. Finally, the dried samples were re-suspended in 0.1 mL of water, 5% formic acid and transferred to glass autosampler vials for LC/MS analysis. Twenty-microliter of extracted supernatant samples was injected into an ultrahigh-performance liquid chromatography (UHPLC) system (Ultimate 3000, Thermo) and run on a positive mode: samples were loaded on to a Reprosil C18 column (2.0mm× 150 mm, 2.5 μm-DrMaisch, Germany) for metabolite separation. For positive ion mode (+) MS analyses, a 0–100% linear gradient of solvent A (ddH2O, 0.1% formic acid) to B (acetonitrile, 0.1%formic acid) was employed over 20 min, returning to 100% A in 2 min and holding solvent A for a 1-min post time hold. Acetonitrile, formic acid, and HPLC-grade water and standards (≥98% chemical purity) were purchased from Sigma Aldrich. Chromatographic separations were made at a column temperature of 30 °C and a flow rate of 0.2 ml/min. The UHPLC system was coupled online with a Q Exactive mass spectrometer (Thermo) scanning in full MS mode (2 μ scans) at resolution of 70,000 in the 67 to 1000 m/z range, a target of 1106 ions and a maximum ion injection time (IT) of 35 ms with 3.8 kV spray voltage, 40 sheath gas and 25 auxiliary gas. Calibration was performed before each analysis against positive or negative ion mode calibration mixes (Pierce, Thermo Fisher, Rockford, IL) to ensure error of the intact mass within the sub ppm range.

Replicates were exported as .mzXML files and processed through MAVEN.8.1. Mass spectrometry chromatograms were created for peak alignment, matching and comparison of parent and fragment ions with tentative metabolite identification (within a 2-p.p.m. mass-deviation range between the observed and expected results against an imported KEGG database). Fold change analysis was performed on the entire metabolomics data set using the MetaboAnalyst 5.0 software. Before the analysis, raw data were normalized by median and autoscaling in order to increase the importance of low-abundance ions without significant amplification of noise. The purpose of fold change (FC) analysis was to compare absolute value change between two group averages and find some features that are changed consistently (i.e. up-regulated or down-regulated) between two groups. In order to analyze a correlation network of the compounds in shared pathways, MetScape, an app implemented in Java and integrated with Cytoscape (version 3.8.2), was used. To further explore the metabolic differences between the two groups of subjects, multivariate statistical analyses were employed on the entire metabolomics data set using the same software. Statistical analyses were employed on the entire metabolomics data set using the same software. The web-based tool MESA or Metabolite Set Enrichment Analysis, which is incorporated into the MetaboAnalyst platform, was used to perform pathway analyses. The Kyoto Encyclopedia of Genes and Genomes (KEGG) human metabolic pathways library was used for pathway analysis.

## Results

Forty-three subjects were enrolled in this observational study and by culturing their PBMC in presence of spike-S1 viral protein, 20 resulted positive to *in vitro* specific IgG secretion (IgG^+^) and 23 were negative (IgG^−^). We use MetaboAnalyst 5.0 platforms to perform untargeted metabolomics to identify in IgG^+^-PBMC possible alterations of relevant metabolites, on the basis that the two groups resulted well clustered by the supervised PLS-DA, as already shown^13^.Fold change analysis was used to identify metabolites altered in IgG^+^-PBMC and involved in defense mechanism against Covid-19, with a FC value greater than 1.5 considered as a significant threshold. The fold change value for each metabolite was Log2 transformed, and the corresponding p-value was −Log10 transformed. With these IgG^+^-PBMC premises, 26 metabolites resulted to have the most significant changes in IgG^+^-PBMC whereas 10 metabolites resulted increased in IgG^−^-PBMC (table and Fig.1).

**Figure 1:**
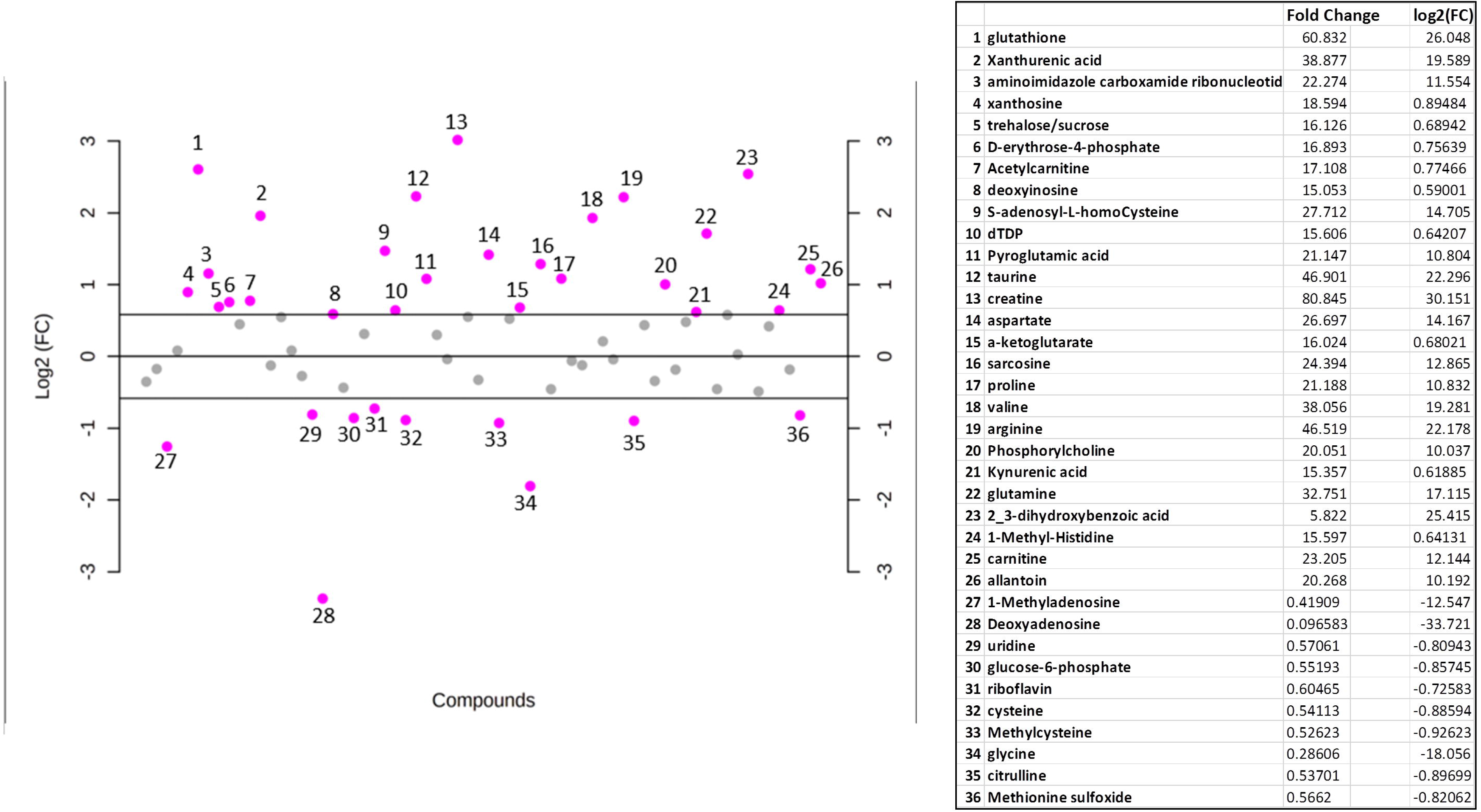
Fold change representation of all analyzed metabolites. Data are log2 fold-change in relative abundance of untargeted metabolites in biological replicates of IgGm+ relative to IgGm-. Dashed lines indicate a log2 fold-change of |1|. Purple dots indicate metabolites with >|1| log2 fold-change and gray dots represent metabolites with <|1| log2 fold-change. Table represents 26 metabolites have shown the most significant changes in the IgGm+ and 10 metabolites resulted increase in IgGm-

The analysis revealed a significant effect of Covid-19 on amino acids metabolism. Namely, gluconeogenic amino acids (e.g., arginine, valine, glutathione, aspartate, proline and glutamine) and taurine (sulfur-containing amino acid) were upregulated in IgG^+^-PBMC. In contrast, cysteine, glycine and methionine sulfoxide (oxidized forms of sulfur-containing amino acid), showed a decrease (Fig. 2).

**Figure 2:**
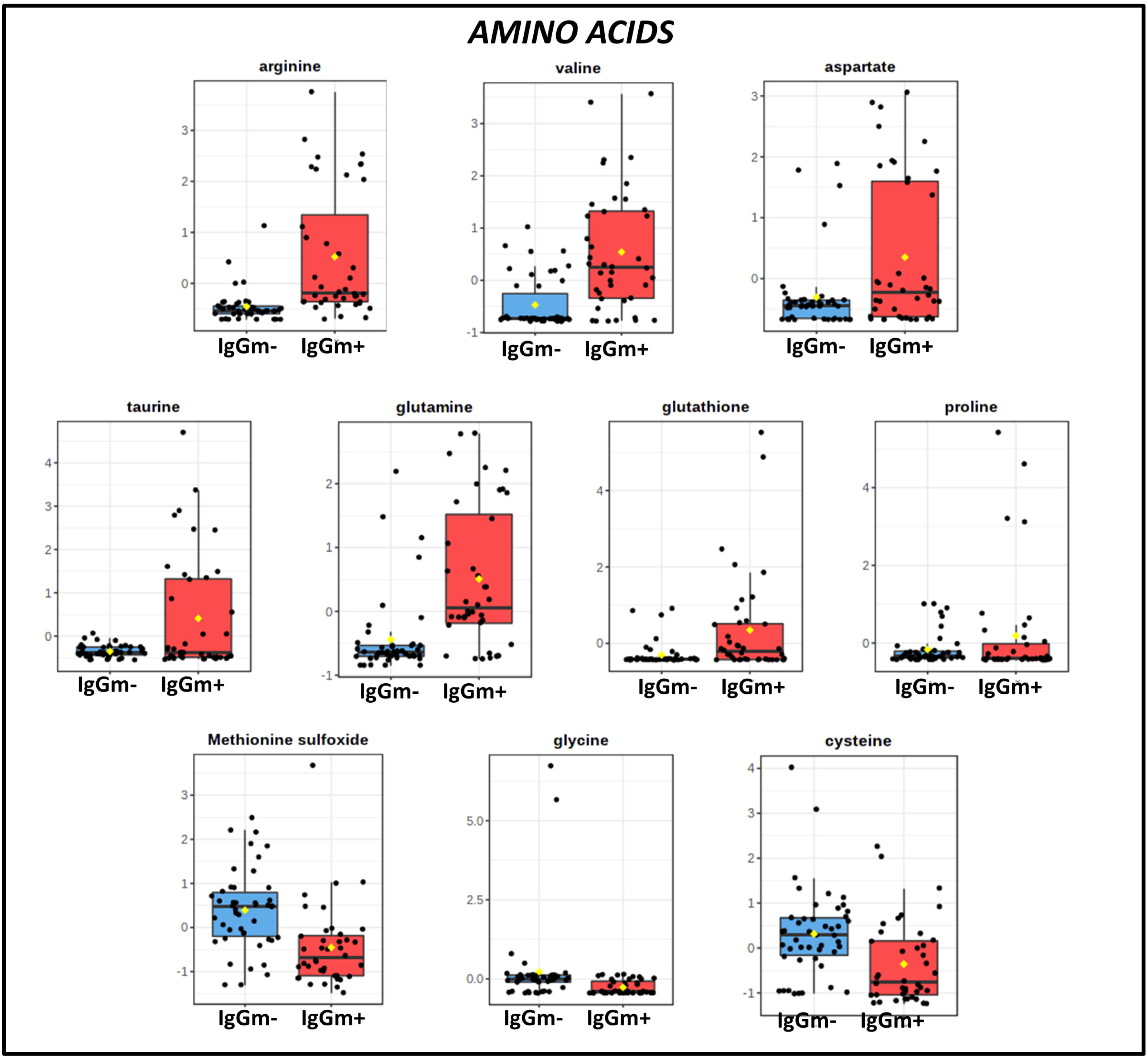
Amino acid levels in PBMC samples.

Based on the online database of metabolic pathways (KEGG, http://www.genome.jp/kegg/), MetScape software running on Cytoscape was used to visualize and interpret metabolites in the context of a global metabolic network (Fig. 3). Network visualizations may be helpful to show connections between metabolites and to understand relationships between compounds and pathways in order to have a high-level overview of changes in metabolic activities caused by Covid-19 in IgG^+^-PBMC. The network reflects a complexity of effects of the pathology and provides further evidence for the involvement of tryptophan metabolism, urea cycle and metabolism of arginine, proline, glutamate, aspartate and asparagine.

**Figure 3:**
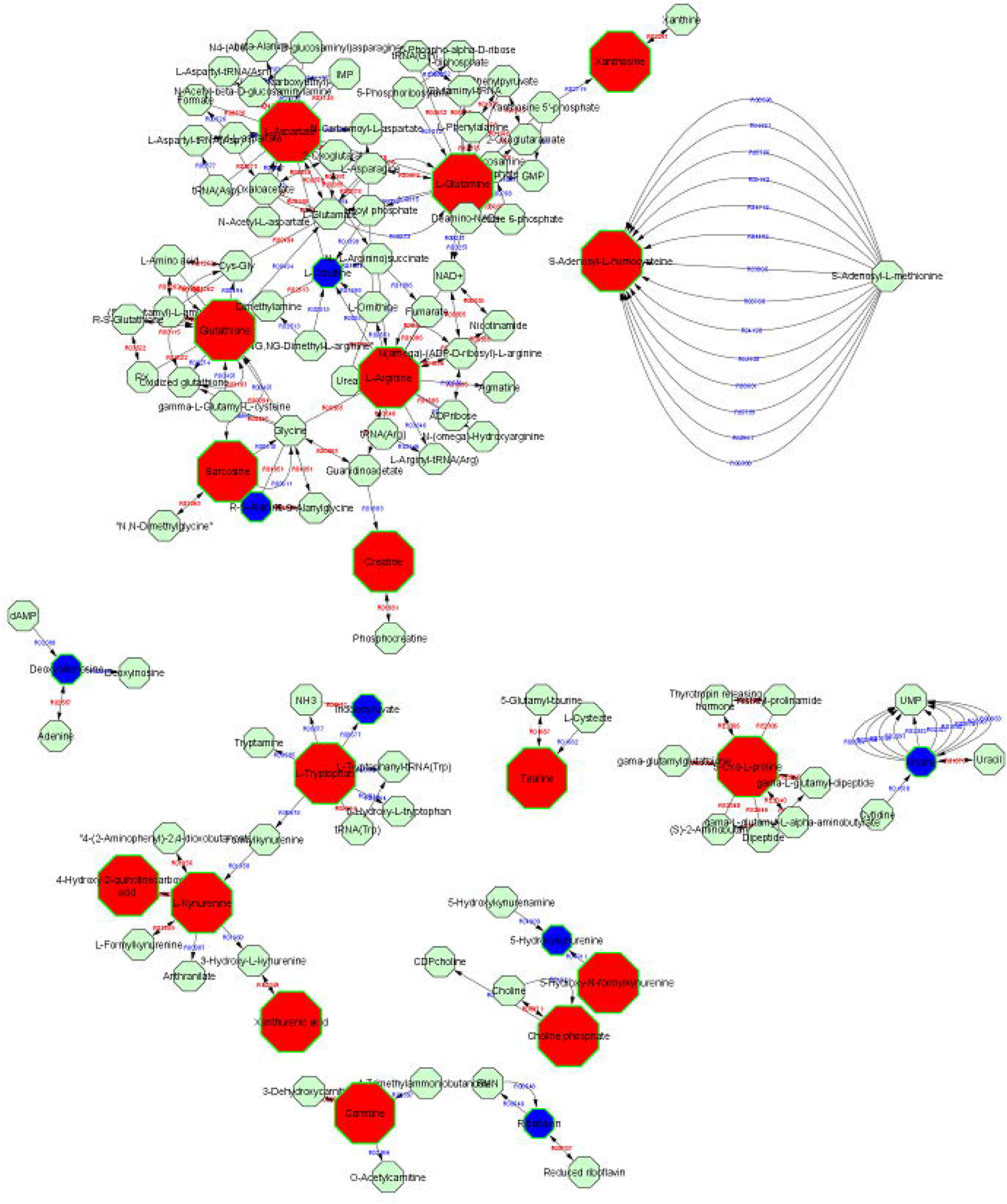
Network analysis by *Cytoscape* including IgGm+ and IgGm- metabolites of VIP score >1. Red metabolites upregulated in IgGm+; blue metabolites down regulated IgGm+ samples; unchanged metabolites, green.

Complementary to the network analysis, Pathway Enrichment Analysis revealed a significant alteration in leukocytes IgGm+ on amino acid metabolism, especially the pathways involving in glycine serine threonine metabolism, arginine biosynthesis, tryptophan metabolism, (Fig. 4). Top hits from these pathways were mapped against the KEGG pathway and tryptophan metabolism and arginine biosynthesis (urea cycle) are highlighted in Figures 4–5.Tryptophan metabolism was the one of top pathways altered in leukocytes displaying an IgG antibody memory to SARS-Cov-2. Specifically, tryptophan increased in the IgGm+ and serotonin did not undergo significant change compared with controls (Fig. 5). At the same time, the reduction in the level of indolepyruvate and the increase in the level of kynurenine, knowing to be rich source for aryl hydrocarbon receptor (AHR) ligands, suggested a restoration of the kynurenine pathway to decrease the cytokines storm (Fig. 5) also confirming by the increased level of taurine in IgGm+.

**Figure 4:**
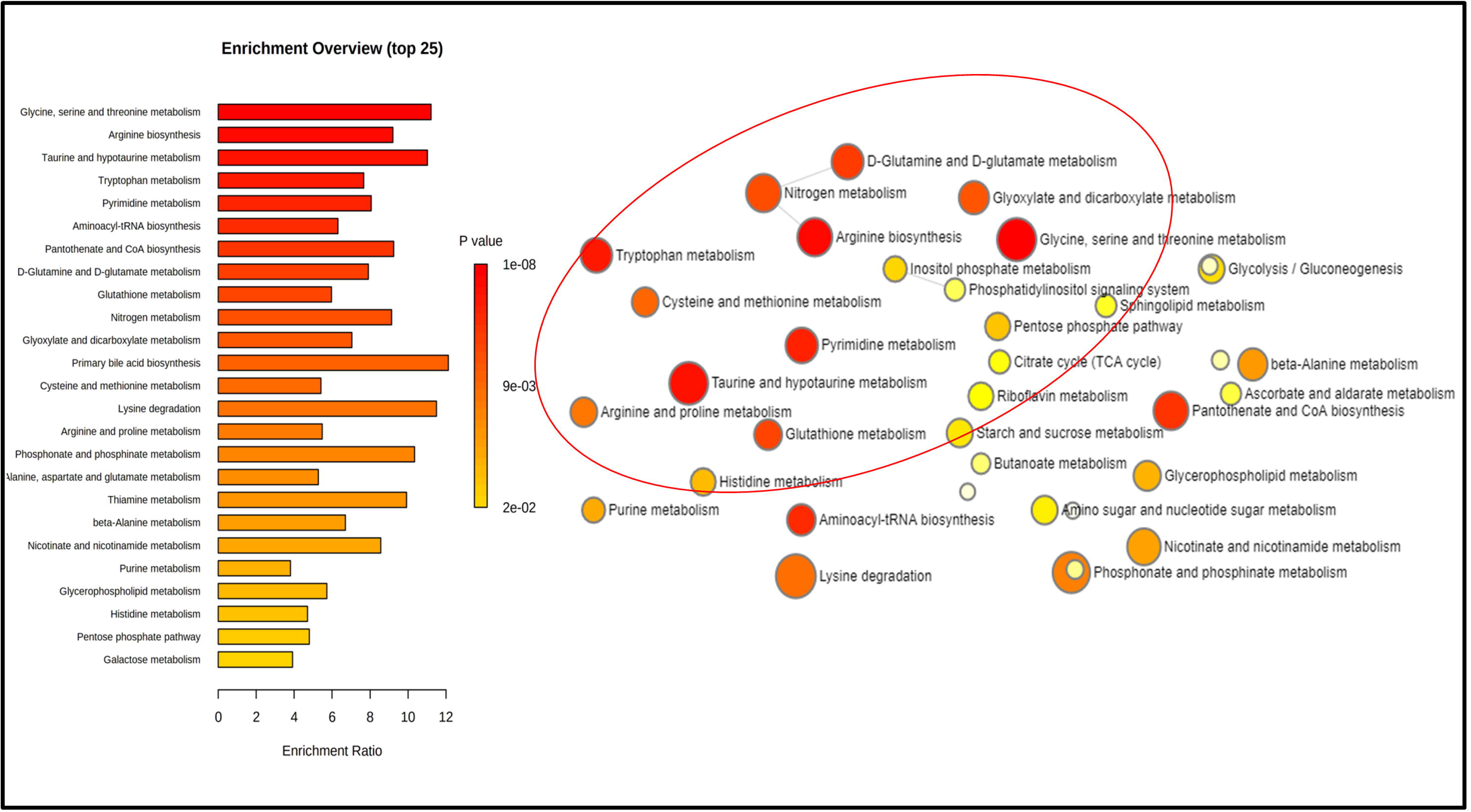
Pathway Enrichment Analysis of untargeted metabolomics. Significant alteration in leukocytes IgGm+ on amino acid metabolism, especially the pathways involved in glycine serine threonine metabolism, arginine biosynthesis, tryptophan metabolism.

**Figure 5:**
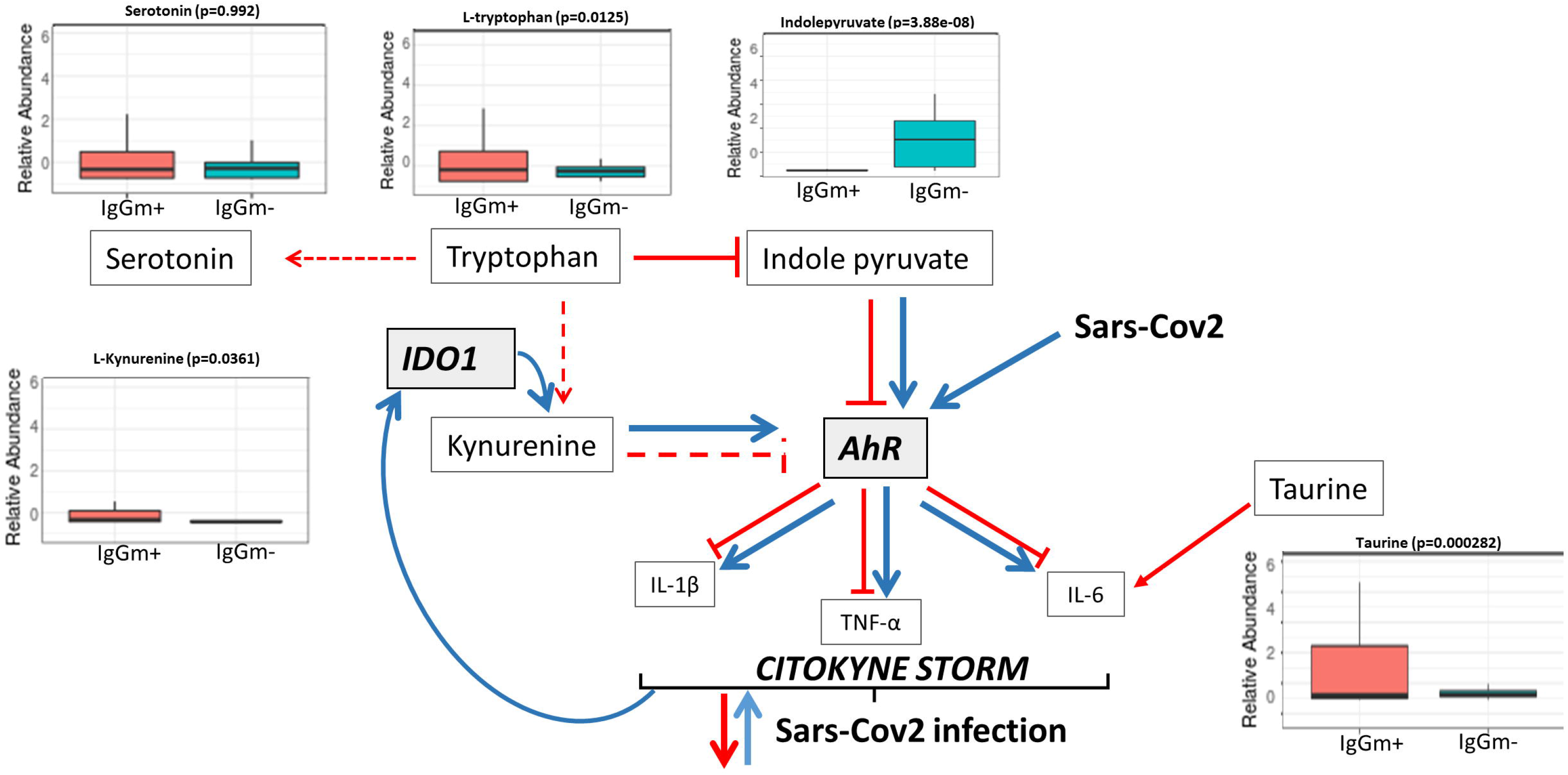
Tryptophan metabolism. Reduction in the level of indolepyruvate and the increase in the level of kynurenine, act on AhR signaling pathways contributing defense against SARS-Cov-2 infection

In addition to Tryptophan metabolism, arginine biosynthesis has a significant impact on leukocytes showing IgG memory. Some metabolites of the Urea cycle (e.g., arginine, aspartate and arginosuccinic acid) were increased in IgGm+ samples, leading to an up regulation of creatine. In contrast, ornithine, glutamic acid and alanine, but not glutamine, did not change in abundance, suggesting an altered homeostasis of the Urea cycle (Fig. 6). As consequence citrulline results fully converted in arginosuccinic acid in IgM+ samples because it wasn’t restored by ornithine and carbamoyl phosphate.

**Figura 6:**
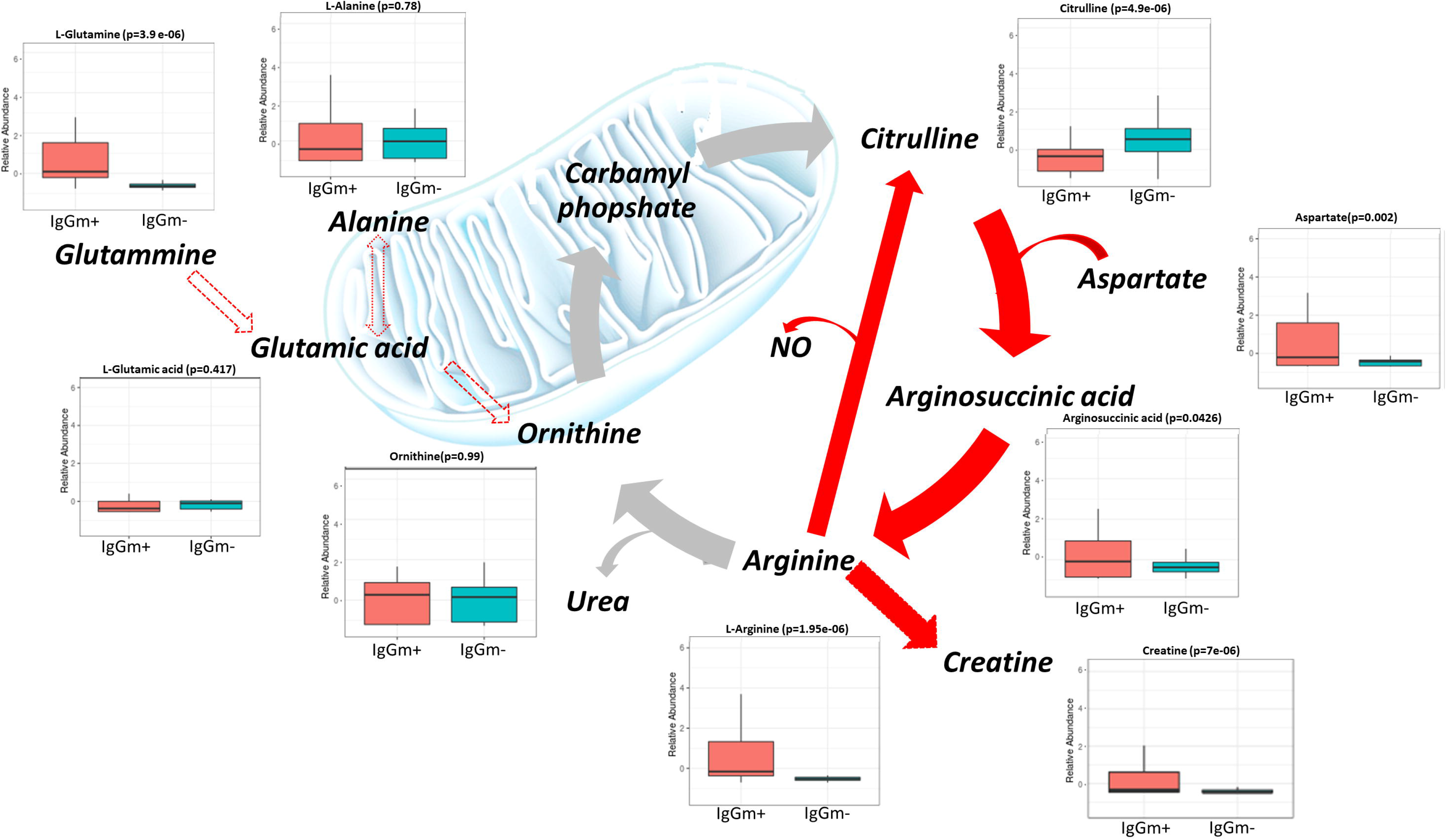
Schematic representation of the urea cycle and its connected metabolism. Urea cycle is associated with the citrulline NO pathway (in Red lines); ornithine, glutamic acid and alanine, but not glutamine, did not change in abundance, suggesting an altered homeostasis of the Urea cycle.

## Discussion

Although host defense mechanisms against SARS-CoV-2 infection are still poorly described, they are critically important in shaping disease course and possible outcome. The untargeted metabolomics profile can integrate the poor knowledge of the molecular mechanisms underlying the infection and reprogramming of leukocyte metabolism after viral pathogenesis. The present study provides the first metabolic characterization of PMCB IgG memory after Sars-coV-2 infection. The results show marked alterations in the metabolism of amino acids in particular in tryptophan and arginine metabolism.

Tryptophan (Trp) metabolism is one of the major pathways affected in IgGm+ leukocytes as shown in our results of non-targeted metabolomic data (Fig. 3). Trp fuels the synthesis of kynurenine (Kyn), serotonin (5-HT) and indoles. A recent study by Gardinassi et al, revealed high involvement of inflammatory networks and increased expression of genes involved in tryptophan metabolism in COVID-19 patients^14^. Interestingly, decrease of tryptophan and the contemporary increase of kynurenine pathway were the major effects of COVID-19 on which regulates inflammation and immunity^7^.This metabolic pathway involves the conversion of tryptophan into L-kynurenine by the enzymes indoleamine 2,3-dioxygenase 1 and 2 (IDO1 and IDO2). IDO enzymes are activated by inflammatory cytokines (IFN-α,-β,-γ, TNF-α) and in particular interleukin-6 (IL-6)^15^. These changes in tryptophan metabolism, due to regulation of IDO enzymes, correlated with a greater increase of interleukin-6 (IL-6) levels^7^, induce a cytokine storm through aryl hydrocarbon receptor (AHR)^16^. The term ‘cytokine storm syndrome’ is perhaps one of the critical hallmarks of COVID-19 disease severity^17^. In our results the increased level of tryptophan, an almost unchanged level of serotonin, and the strongly decreased level of indole pyruvate support the hypothesis of the restoration of kynurenine pathway through attenuation of IDO activity. In addition to kynurenine, also indole pyruvate are endogenous ligands that activate AhR^18,19^. Therefore the reduction level of indole pyruvate negatively modulates the AHR activity in leukocytes which display IgG memory for spike-S1 antigen 60-90 days after infection. Moreover the increased level of Taurine in IgGm+ attenuated IL-6 level^20^ cooperating to reduce inflammation. Increasing evidence from several studies show that tryptophan can reduce inflammatory reactions and enhance the immune system^21^. Furthermore, recent studies have suggested a link between tocilizumab immunosuppressive therapy and tryptophan metabolism in COVID-1. In fact, tocilizumab has been proposed as a drug to counteract hyperinflammatory responses in ICU patients with COVID-19^22^. This possibility indicates that tryptophan-rich sources could be beneficial for COVID-19 subjects ^23^. The potential relevance of these observations confirm the impact of Trp and its metabolites against the severity of COVID-19, and our results highlighted for the first time to our knowledgement the biochemical mechanism adopted by leukocytes to counteract viral infection by reactivating the tryptophan metabolism.

Another metabolism significantly influenced by Sars-Cov2 virus was arginine biosynthesis. Arginine is a semi-essential amino acid that can be obtained from the diet or produced in certain cells via the complete or partial urea cycle. In the urea cycle, when arginine is catabolized by arginase (Arg1), the products are urea, ornithine, polyamines and proline, and when degraded by NO, the products are a large amount of NO and citrulline^24^. The ornithine used for citrulline production can originate from alanine and glutamate/glutamine reactions. In our results arginine wasn’t converted to ornithine. In fact this latter was not fuel by glutamine that are major substrates for citrulline production^25^. At the same time the increased level of glutamine is in agreement with the role of this amino acid in boosting the immune system especially by inhibition of inflammatory responses^26^. Indeed, adding enteral glutamine to the normal nutrition in the early period of Covid-19 infection may lead to a shortened hospital stay and lead to less need for ICU^27^.

The conversion of arginine in citrulline rather than ornithine leads to a higher release of NO. In the immune system, the NO produced in macrophages and neutrophils is necessary to kill invasive microorganisms (such as viruses, bacteria, and fungi) and activate the immune cells in defense mechanisms^28,29^. Arginine is a substrate for nitric oxide (NO) production, which can induce antiviral activity against RNA viruses, such as SARS-CoV-2^29^. The accumulation of this amino acid in IgGm+^13^ and consequently alteration of urea cycle was probably due to an restoration of arginase enzyme (Arg1). It has been reported that the Arg1 gene is located in the cytoplasm and is strongly expressed in the liver but, in addition to its metabolic role in the hepatic urea cycle, it can regulate immune responses. In fact, in humans, arginase was detected in the peripheral blood mononuclear cell (PBMC)^31^ and several studies highlighted that Arg1 inhibits immunity against intracellular pathogens and represses T-cell-mediated inflammatory damage^32,33^. Arg1 up-regulation might be associated with higher virus load in COVID-19 patients^34^. Since Arg1 can limit the bioavailability of L-arginine, the inhibition of Arg1 can drive the recycling of L-citrulline to generate L-arginine for the production of NO, paving the way for developing antiviral immunity in IgGm+.

A significant proportion of L-arginine flux is attributable to synthesis of creatine by the enzyme L-arginine:glycine amidinotransferase (AGAT). Generation of creatine is estimated to consume ~70% of labile methyl groups, with S-adenosyl-methionine (SAM) serving as the methyl donor^35^. This methylation demand may reduce the available methyl which therefore leads SAM to be metabolized into S-adenosyl homocysteine (SAH) and finally into homocysteine and indirectly limits methylation of the 5’ cap of viral messenger RNA of coronavirus^13^. So it is not surprising that the Creatine Kinase (enzyme that catalyses the conversion of creatine to create phosphocreatine) levels are higher in COVID-19 patients and are associated with a more severe prognosis in COVID-19^36^.

In conclusion, metabolomics analysis determines the developmental, physiological, and pathological states of a biological system thereby representing a powerful tool for precision medicine. In this study, for the first time, we provide novel insights into later stages of immune defense against SARS-Cov-2 infection, namely when circulating antibodies may be absent, but an antibody memory is present. After viral pathogenesis, metabolomics analysis reveals a reprogrammed of leukocyte metabolism through activation of specific amino acid pathways related to protective immunity against SARS-CoV-2.

## Acknowledgments

This work was supported by PRIN 2017 from University of Viterbo. and by Ministero dell’Istruzione, dell’Università e della Ricerca (MIUR), Rome, Italy.

## References

1. Baj J, Karakuła-Juchnowicz H, Teresiński G, Buszewicz G, Ciesielka M, Sitarz E, et al. COVID-19: Specific and Non-Specific Clinical Manifestations and Symptoms: The Current State of Knowledge. J Clin Med. 2020 Jun 5;9(6):1753. doi: 10.3390/jcm9061753.

2. Gautam S, Gautam A, Chhetri S, Bhattarai U. Immunity Against COVID-19: Potential Role of Ayush Kwath. J Ayurveda Integr Med. 2020 Aug 17. doi: 10.1016/j.jaim.2020.08.003.

3. Turski WA, Wnorowski A, Turski GN, Turski CA, Turski L. AhR and IDO1 in pathogenesis of Covid-19 and the “Systemic AhR Activation Syndrome:” a translational review and therapeutic perspectives. Restor Neurol Neurosci. 2020; 38(4):343–354. doi:10.3233/RNN-201042.

4. Tay MZ, Poh CM, Rénia L, MacAry PA, Ng LFP. The trinity of COVID-19: immunity, inflammation and intervention. Nat Rev Immunol. 2020 Jun;20(6):363–374. doi: 10.1038/s41577-020-0311-8.

5. Shah VK, Firmal P, Alam A, Ganguly D, Chattopadhyay S. Overview of Immune Response During SARS-CoV-2 Infection: Lessons From the Past. Front Immunol. 2020 Aug 7;11:1949. doi: 10.3389/fimmu.2020.01949.

6. Stockinger B, Bourgeois C, Kassiotis G. CD4+ memory T cells: functional differentiation and homeostasis. Immunol Rev. 2006 Jun;211:39–48. doi: 10.1111/j.0105-2896.2006.00381.x.

7. Thomas T, Stefanoni D, Reisz JA, Nemkov T, Bertolone L, Francis RO, Hudson KE, et al. COVID-19 infection alters kynurenine and fatty acid metabolism, correlating with IL-6 levels and renal status. JCI Insight. 2020 Jul 23;5(14):e140327. doi: 10.1172/jci.insight.140327.

8. Danlos FX, Grajeda-Iglesias C, Durand S, et al. Metabolomic analyses of COVID-19 patients unravel stage-dependent and prognostic biomarkers. Cell Death & Disease. 2021 Mar;12(3):258. DOI: 10.1038/s41419-021-03540-y.

9. Roberts I, Wright Muelas M, M. Taylor J, S. Davison A, Xu Y, M. Grixti J, et al. Untargeted metabolomics of COVID-19 patient serum reveals potential prognostic markers of both severity and outcome. medRxiv (2020).12.09.20246389; doi: https://doi.org/10.1101/2020.12.09.20246389.

10. Dan JM, Mateus J, Kato Y, Hastie KM, Yu ED, Faliti CE, Grifoni A, et al. Immunological memory to SARS-CoV-2 assessed for up to 8 months after infection. Science. 2021 Feb 5;371(6529):eabf4063. doi: 10.1126/science.abf4063.

11. Mahase, E. Covid-19: What do we know about “long covid”? BMJ 2020 Jul 14;370:m2815. doi: 10.1136/bmj.m2815.

12. Zarletti G, Tiberi M, De Molfetta V, Bossù M, Toppi E, Bossù P, Scapigliati G. A Cell-Based ELISA to Improve the Serological Analysis of Anti-SARS-CoV-2 IgG. Viruses. 2020; 12(11):1274. https://doi.org/10.3390/v12111274

13. Fanelli G, Gevi F, Zarletti G, Tiberi M, De Molfetta V, Scapigliati G, Timperio A.M.. An altered metabolism in leukocytes showing in vitro igG memory from SARS-CoV-2-infected patients. bioRxiv 2021.05.27.445918; doi: https://doi.org/10.1101/2021.05.27.445918.

14. Gardinassi LG, Souza COS, Sales-Campos H, Fonseca SG. Immune and Metabolic Signatures of COVID-19 Revealed by Transcriptomics Data Reuse. Front Immunol. 2020 Jun 26;11:1636. doi: 10.3389/fimmu.2020.01636.

15. Kim S, Miller BJ, Stefanek ME, Miller AH. Inflammation-induced activation of the indoleamine 2,3-dioxygenase pathway: Relevance to cancer-related fatigue. Cancer. 2015 Jul 1;121(13):2129–36. doi: 10.1002/cncr.29302.

16. Anderson G, Carbone A, Mazzoccoli G. Tryptophan Metabolites and Aryl Hydrocarbon Receptor in Severe Acute Respiratory Syndrome, Coronavirus-2 (SARS-CoV-2) Pathophysiology. Int J Mol Sci. 2021 Feb 5;22(4):1597. doi: 10.3390/ijms22041597.

17. Fara A, Mitrev Z, Rosalia RA, Assas BM. Cytokine storm and COVID-19: a chronicle of pro-inflammatory cytokines. Open Biol. 2020 Sep;10(9):200160. doi: 10.1098/rsob.200160.

18. Chowdhury G, Dostalek M, Hsu EL, Nguyen LP, Stec DF, Bradfield CA, Guengerich FP. Structural identification of Diindole agonists of the aryl hydrocarbon receptor derived from degradation of indole-3-pyruvic acid. Chem Res Toxicol. 2009 Dec;22(12):1905–12. doi: 10.1021/tx9000418.

19. Aoki R, Aoki-Yoshida A, Suzuki C, Takayama Y. Indole-3-Pyruvic Acid, an Aryl Hydrocarbon Receptor Activator, Suppresses Experimental Colitis in Mice. J Immunol. 2018 Dec 15;201(12):3683–3693. doi: 10.4049/jimmunol.1701734.

20. Iwegbulem O, Wang J, Pfirrmann RW, Redmond HP. The role of taurine derivatives in the putative therapy of COVID-19-induced inflammation. Ir J Med Sci. 2021 Feb 18:1–2. doi: 10.1007/s11845-021-02522-5.

21. Essa MM, Hamdan H, Chidambaram SB, Al-Balushi B, Guillemin GJ, Ojcius DM, Qoronfleh MW. Possible role of tryptophan and melatonin in COVID-19. Int J Tryptophan Res. 2020 Aug 21;13:1178646920951832. doi: 10.1177/1178646920951832.

22. Belladonna ML, Orabona C. Potential Benefits of Tryptophan Metabolism to the Efficacy of Tocilizumab in COVID-19. Front Pharmacol. 2020 Jun 19;11:959. doi: 10.3389/fphar.2020.00959.

23. Shader RI. COVID-19, interferons, and depression: A commentary. Psychiatry Research. 2020 Sep;291:113198. DOI: 10.1016/j.psychres.2020.113198.

24. Wu G, Bazer FW, Davis TA, Kim SW, Li P, Marc Rhoads J, Carey Satterfield M, Smith SB, Spencer TE, Yin Y. Arginine metabolism and nutrition in growth, health and disease. Amino Acids. 2009 May;37(1):153–68. doi: 10.1007/s00726-008-0210-y.

25. Marini JC. Interrelationships between glutamine and citrulline metabolism. Curr Opin Clin Nutr Metab Care. 2016 Jan;19(1):62–6. doi: 10.1097/MCO.0000000000000233

26. de Oliveira DC, da Silva Lima F, Sartori T. et al. Glutamine metabolism and its effects on immune response: molecular mechanism and gene expression. Nutrire 41, 14 (2016). https://doi.org/10.1186/s41110-016-0016-8

27. Cengiz M, Borku Uysal B, Ikitimur H, Ozcan E, Islamoğlu MS, Aktepe E, Yavuzer H, Yavuzer S. Effect of oral l-Glutamine supplementation on Covid-19 treatment. Clin Nutr Exp. 2020 Oct;33:24–31. doi: 10.1016/j.yclnex.2020.07.003.

28. Lisi F, Zelikin AN, Chandrawati R. Nitric Oxide to Fight Viral Infections. Adv. Sci., 2021. p. 2003895. https://doi.org/10.1002/advs.202003895

29. Wink DA, Hines HB, Cheng RY, Switzer CH, Flores-Santana W, Vitek MP, Ridnour LA, Colton CA. Nitric oxide and redox mechanisms in the immune response. J Leukoc Biol. 2011 Jun;89(6):873–91. doi: 10.1189/jlb.1010550. Epub 2011 Jan 13. PMID: 21233414; PMCID: PMC3100761.

30. Derakhshani A, Hemmat N, Asadzadeh Z, Ghaseminia M, Shadbad M.A, Jadideslam, G, et al. Arginase 1 (Arg1) as an Up-Regulated Gene in COVID-19 Patients: A Promising Marker in COVID-19 Immunopathy. J. Clin. Med. 2021, 10, 1051. https://doi.org/10.3390/jcm10051051.

31. Ochoa JB, Bernard AC, O’Brien WE, Griffen MM, Maley ME, Rockich AK, et al. Arginase I expression and activity in human mononuclear cells after injury. Ann Surg. 2001 Mar;233(3):393–9. doi: 10.1097/00000658-200103000-00014.

32. Zea AH, Rodriguez PC, Atkins MB, Hernandez C, Signoretti S, Zabaleta J, et al. Arginase-producing myeloid suppressor cells in renal cell carcinoma patients: a mechanism of tumor evasion. Cancer Res. 2005 Apr 15;65(8):3044–8. doi: 10.1158/0008-5472.

33. Pesce JT, Ramalingam TR, Mentink-Kane MM, Wilson MS, El Kasmi KC, Smith AM, et al. Arginase-1-expressing macrophages suppress Th2 cytokine-driven inflammation and fibrosis. PLoS Pathog. 2009 Apr;5(4):e1000371. doi: 10.1371/journal.ppat.1000371.

34. Hemmat N, Derakhshani A, Bannazadeh Baghi H, Silvestris N, Baradaran, & De Summa S. Neutrophils, Crucial, or Harmful Immune Cells Involved in Coronavirus Infection: A Bioinformatics Study. Frontiers in genetics, 2020 11, 641. https://doi.org/10.3389/fgene.2020.00641

35. Jahangir E, Vita JA, Handy D, Holbrook M, Palmisano J, Beal R, Loscalzo J, Eberhardt RT. The effect of L-arginine and creatine on vascular function and homocysteine metabolism. Vasc Med. 2009 Aug;14(3):239–48. doi: 10.1177/1358863X08100834.

36. Orsucci D, Trezzi M, Anichini R, Blanc P, Barontini L, Biagini C, et al. Increased Creatine Kinase May Predict A Worse COVID-19 Outcome. J Clin Med. 2021 Apr 16;10(8):1734. doi: 10.3390/jcm10081734.

