## supplementary for "An untargeted metabolomic approach to identify antiviral defense mechanisms in memory leukocytes secreting *in vitro* IgG anti-SARS-Cov-2"

| Age/Gender | *in vitro* IgG memory (Cell-ELISA)* |
|---|---|
| 33 F | 0,039 |
| 33 F | **0,239** |
| 33 M | 0,023 |
| 35 M | 0,02 |
| 36 M | **0,07** |
| 36 F | **0,292** |
| 39 F | **0,223** |
| 40 F | **0,291** |
| 40 M | 0,019 |
| 40 M | 0,026 |
| 42 F | **0,258** |
| 42 F | **0,087** |
| 42 M | 0,024 |
| 43 M | 0,024 |
| 44 M | 0,023 |
| 44 M | 0,02 |
| 45 M | 0,023 |
| 45 M | 0,017 |
| 46 M | 0,058 |
| 47 F | **0,293** |
| 47 F | **0,085** |
| 49 M | 0,021 |
| 49 M | 0,022 |
| 49M | 0,025 |
| 50 M | 0,019 |
| 50 F | **0,141** |
| 53 M | 0,067 |
| 54 M | 0,014 |
| 57 F | **0,177** |
| 58 F | **0,245** |
| 63 M | 0,038 |
| 67 F | **0,285** |
| 68 F | 0,05 |
| 70 F | **0,118** |
| 71 M | **0,082** |
| 72 M | 0,023 |
| 72 M | 0,033 |
| 72 F | 0,027 |

| | |
|---|---|
| 72 F | **0,116** |
| 73 M | 0,037 |
| 74 F | **0,217** |
